# Critical genetic program for *Drosophila* imaginal disc regeneration revealed by single-cell analysis

**DOI:** 10.1101/2021.07.08.451678

**Authors:** Melanie I. Worley, Nicholas J. Everetts, Riku Yasutomi, Nir Yosef, Iswar K. Hariharan

## Abstract

Whether regeneration is primarily accomplished by re-activating gene regulatory networks used previously during development or by activating novel regeneration-specific transcriptional programs remains a longstanding question. Currently, most genes implicated in regeneration also function during development. Using single-cell transcriptomics in regenerating *Drosophila* wing discs, we identified two regeneration-specific cell populations within the blastema. They are each composed of cells that upregulate multiple genes encoding secreted proteins that promote regeneration. In this regenerative secretory zone, the transcription factor Ets21C controls the expression of multiple regeneration-promoting genes. While eliminating *Ets21C* function has no discernible effect on development, it severely compromises regeneration. This Ets21C-dependent gene regulatory network is also activated in blastema-like cells in tumorous discs, suggesting that pro-regenerative mechanisms can be co-opted by tumors to promote aberrant growth.

---

A long-standing question in the field of regenerative biology is whether regeneration is mainly accomplished by reactivation of gene regulatory networks (GRNs) used during earlier stages of development or, alternatively, by GRNs that are specifically activated during regeneration. Studies of regenerating tissues have provided evidence for cellular states that are not observed during normal development and for patterns of gene expression that seem specific for regeneration (for example Gerber *et al*., 2018; Aztekin *et al*., 2019). However, so far, there is little evidence for genes that are needed for regeneration but not for normal development.

To identify transcriptional programs initiated during regeneration, we examined regeneration of *Drosophila* larval wing imaginal discs, the epithelial tissues that differentiate into the adult wings and thorax. Imaginal discs are capable of regenerating after damage through the formation of a blastema, defined by localized proliferation and increased cellular plasticity (reviewed by Worley *et al*., 2012). To search for regeneration-specific GRNs, we compared regenerating and developing wing discs using single-cell transcriptomics. Tissue damage was induced by temporarily expressing the pro-apoptotic TNF ortholog *eiger* within the wing pouch, the portion of the disc that generates the wing blade (Smith-Bolton *et al*., 2009) (**supplemental fig. 1**). Subsequent regeneration occurs by localized cell proliferation and cell-fate re-specification. We collected wing discs after 24 hours of regeneration, approximately one third of the way through the regenerative process, and sequenced a total of 14,320 cells from two biological replicates, with an average of >3,000 genes detected per cell. Three major cell types were identified: epithelial cells, myoblasts, and hemocytes (**supplemental fig. 2**). Since imaginal disc regeneration is driven by epithelial cell proliferation (Smith-Bolton *et al*., 2009; Worley *et al*., 2012), we focused further analysis on these cells.

To identify potential regeneration-specific GRNs, we harmonized data from epithelial cells from regenerating discs with our previously collected data from undamaged discs (Everetts *et al*., 2021) using scVI (Gayoso *et al*., 2021) (see Materials and Methods) (**Figure 1A, B**). We assigned cell clusters to specific subregions of the wing disc epithelium based on the expression of known marker genes (Bageritz *et al*., 2019; Deng *et al*., 2019; Zappia *et al*., 2020; Everetts *et al*., 2021) (**Figure 1B, C; supplemental fig. 3**). As expected, cell clusters with pouch identity were underrepresented in the regenerating sample, as this portion of the tissue was ablated (**supplemental fig. 3**). From our single-cell analysis, we observed two clusters, denoted Blastema1 and Blastema2, that were almost exclusively composed of cells from the regenerating sample (181/186 and 519/564 cells, respectively) (**Figure 1B; supplemental fig. 3**). Within these two regeneration-specific clusters, we observed the upregulation of genes known to be induced around the site of damage, including the Wnt ligands *wingless (wg)* and *Wnt6* (Harris *et al*., 2016), *Matrix metalloproteinase 1 (Mmp1)*, and *Insulin-like peptide 8 (Ilp8)* (**Figure 1C, D**).

**Figure 1.**
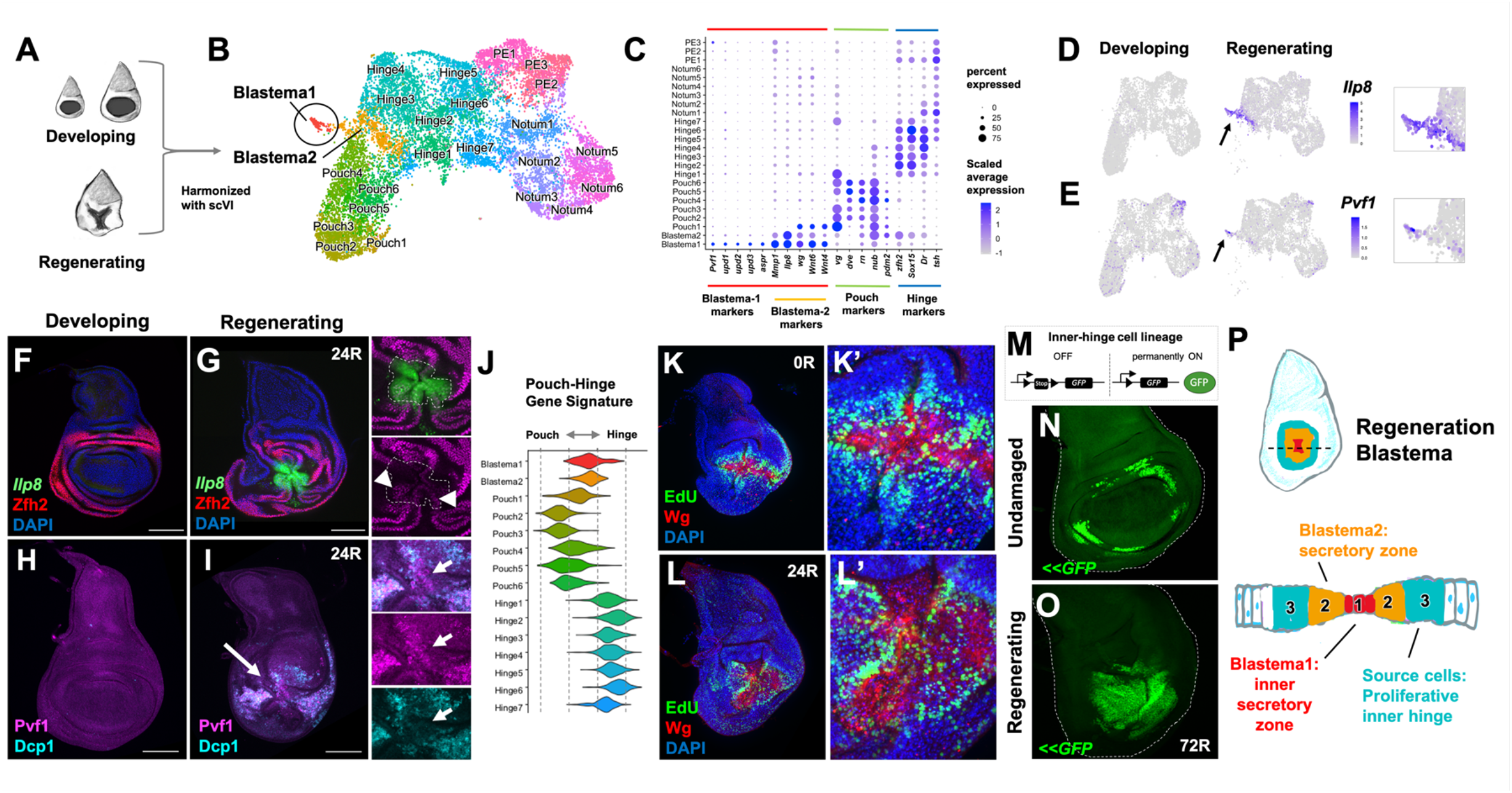
Single-cell analysis reveals two distinct cell states in the regeneration blastema. (**A**) Diagram of imaginal disc samples compared by scRNAseq. (**B**) UMAP of harmonized epithelial cells. (**C**) Dot plot summarizing gene expression for cluster marker genes. (**D, E**) Expression of *Ilp8* (**D**) and *Pvf1* (**E**) as visualized on UMAP. (**F-I**) Developing and regenerating wing discs, after 24 h of regeneration (24R), (**F, G)** with an *Ilp8-GFP* reporter stained with anti-Zfh2 (hinge marker), and (**H, I**) stained with anti-Pvf1 and anti-cleaved Death caspase-1 (Dcp-1) (detects apoptotic cells and debris). (**J**) Pouch-Hinge gene signature analysis of blastema cell clusters. (**K, L**) Regenerating wing discs at 0R and 24R with cells in S-phase visualized by EdU incorporation. (**M**) Schematic of lineage-tracing technique with an inner-hinge enhancer. Lineage tracing in (**N**) normal development and (**O**) following 72h or regeneration. (**P**) Schematic of distinct cell types of the blastema.

Both Blastema1 and Blastema2 clusters express *Ilp8*, which is strongly upregulated around the site of damage in the regenerating disc (**Figure 1F, G**). However, Blastema2 showed a higher expression of hinge-identity markers, such as transcription factor *Zn finger homeodomain 2 (zfh2)*, than Blastema1 (**Figure 1C**), suggesting that these cells might occupy a more proximal (outer) position. Indeed, in regenerating tissue, we observed higher Zfh2 expression in the outer ring of *Ilp8*-expressing cells (**Figure 1G**). In contrast, Blastema1 cells expressed higher levels of the *unpaired (upd1, upd2, upd3)* ligands, *asperous (aspr)*, and *PDGF- and VEGF-related factor 1 (Pvf1)* (**Figure 1C, E**). The Upd ligands activate the JAK/STAT pathway, which is important for cellular plasticity and regeneration (Katsuyama *et al*., 2015; Santabarbara-Ruiz *et al*., 2015; La Fortezza *et al*., 2016; Worley *et al*., 2018). The gene *aspr* encodes a secreted protein with multiple EGF-repeats important for regeneration (Harris *et al*., 2020). Pvf1 binds to its receptor Pvr and the resulting signaling is known to contribute to wound healing (Wu *et al*., 2009), and homologs are involved in regeneration in other systems (Currie *et al*., 2016; Johnson *et al*., 2020). We determined that *Pvf1, upd3*, and *Ilp8* were all expressed at the center of the blastema **(Figure 1H, I; supplemental fig. 4**), which is surrounded by cells that express *Ilp8* but not *Pvf1* or *upd3*. Thus, the Blastema1 cells are located at the center of the blastema and are surrounded by Blastema2 cells; cells in both regions secrete ligands, some of which are known to promote regeneration, and are likely acting on the surrounding tissue. We refer to these regions together as the regenerative secretory zone.

From our single-cell analysis, we used gene signatures to determine that the cells within the regenerative secretory zone were in an intermediate state between hinge and pouch identities (**Figure 1J; supplemental fig. 5**). This finding suggested that these cells were derived from the surrounding inner-hinge region and were in the process of acquiring more distal pouch fates. To investigate this process, we examined the location of proliferating cells and found high levels of EdU incorporation surrounding the regenerative secretory zone (**Figure 1K**) (Cosolo *et al*., 2019). As regeneration proceeded, the EdU incorporation extended more centrally to occur within the regenerative secretory zone (**Figure 1L**). To determine if these proliferating cells are reprogrammed during regeneration to replace the ablated pouch, we performed a lineage-tracing experiment with an enhancer that is normally only expressed in a ring of cells of the inner-hinge. In the absence of pouch ablation, these cells and their progeny remain confined to the hinge (**Figure 1M, N**). However, after regeneration following pouch ablation, most of the regenerated pouch was derived from cells that once expressed this enhancer (**Figure 1O**). Thus, the ablated pouch is regenerated by the proliferation and reprogramming of more proximally fated inner hinge cells, likely driven by the ligands secreted by the regenerative secretory zone (**Figure 1P**).

To search for a regulator of these regeneration-specific transcriptional changes, we analyzed our single-cell data for a transcription factor that was specifically expressed within the blastema cells. We found that *Ets at 21C (Ets21C)* was specifically expressed during regeneration, primarily within the cells of the regenerative secretory zone, and not in cells from developing wing discs which were undamaged (**Figure 2A-C**). *Ets21C* was also upregulated after physically wounding of the wing disc (**Figure 2D**), implying that *Ets21C* is involved in a general regeneration response. *Ets21C* had previously been shown to be upregulated during disc regeneration by bulk sequencing of blastema-enriched cells (Khan *et al*., 2017). Our single-cell data indicates that *Ets21C* expression is highly correlated with *Ilp8* and *Mmp1* expression during regeneration (**supplemental fig. 6**). *Ets21C* expression was induced during the genetic ablation period and was maintained throughout regeneration (**supplemental fig. 7**), suggesting that Ets21C could function at multiple stages of regeneration.

**Figure 2.**
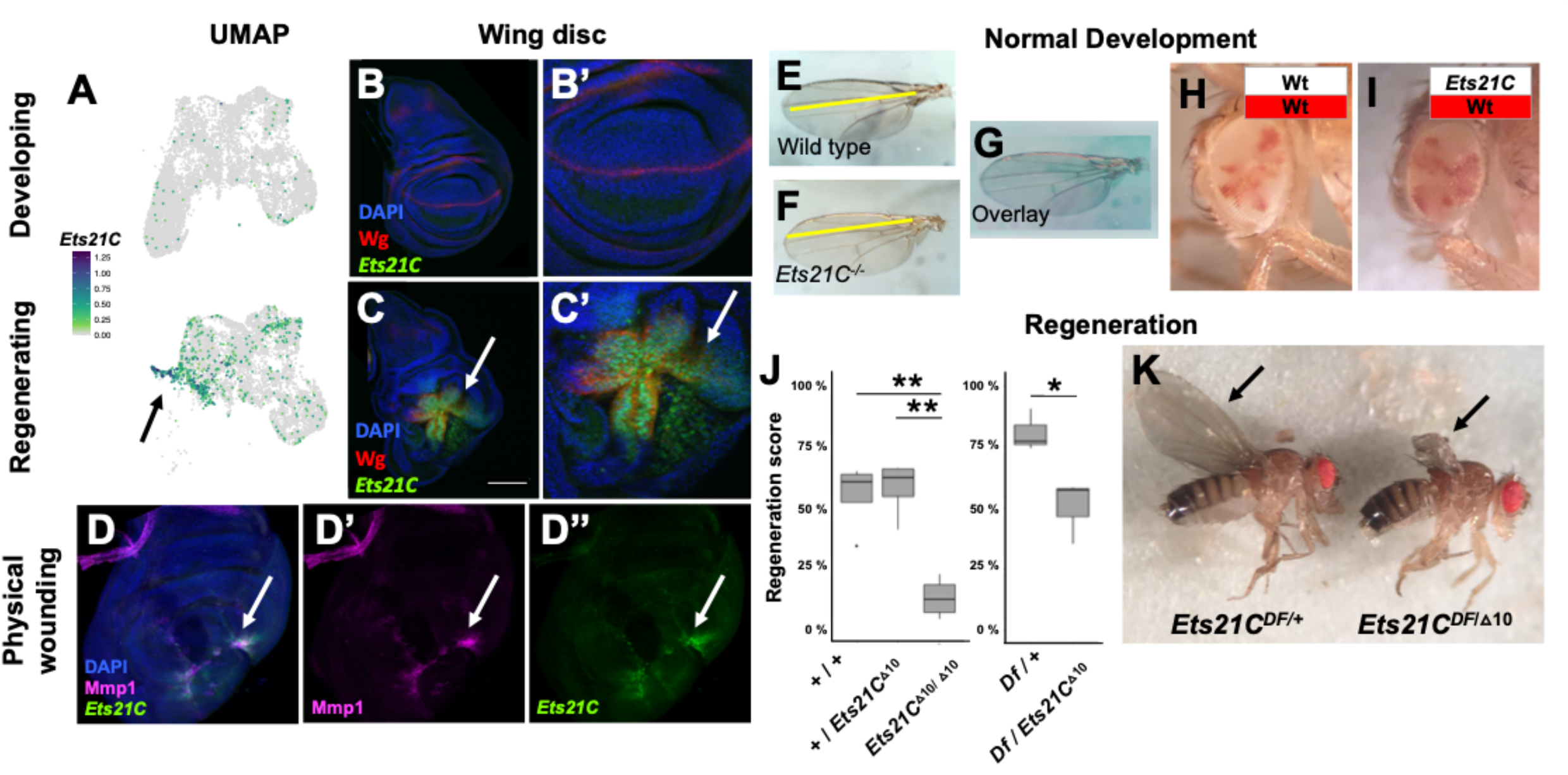
Transcription factor Ets21C is specifically required for regeneration. (**A**) *Ets21C* expression in developing and regenerating scRNAseq data. (**B, C**) *Ets21C-GFP* expression in (**B**) developing and (**C**) regenerating wing discs. (**D**) *Ets21C-GFP* expression following physically wounding disc through larval cuticle. (**E-G**) Wing blades from wild-type and *Ets21C* mutant animals, raised in standard conditions, and shown overlaid. (**H, I**) Mosaic adult eyes with control (**H**) and *Ets21C* (**I**) cells marked by the absence of red pigment. Note that *Ets21C* mutant cells (**I**) contribute to tissue at a similar proportion as control cells (**H**). (**J, K**) The extent of regeneration, as scored by the size of the resulting wing blades, p values: * <0.05, ** <0.005 (see Materials and Methods).

To determine if *Ets21C* was important either for normal development or for regeneration, we turned to mutant analysis. First, we observed that homozygous *Ets21C^-/-^* null mutants generate viable and fertile adults, as previously noted (Mundorf *et al*., 2019), whose wings were of normal size and shape (**Figure 2E-G**). By generating mosaic eyes composed of marked wild-type cells and *Ets21C^-/-^* mutant cells, we found that mutant cells did not display defects in cell proliferation even in a competition scenario with wild-type cells (**Figure 2H, I**). Thus, *Ets21C* is dispensable for normal development and its absence does not impair cell proliferation.

Next, we tested if *Ets21C* is essential for imaginal disc regeneration. Following our genetic ablation assay, homozygous null *Ets21C^-/-^* mutants showed a dramatic defect in the extent of wing regeneration when compared to either *Ets21C^+/-^* heterozygotes or wild-type controls (**Figure 2J**). This effect was observed with the null mutation in trans to a chromosomal deletion (**Figure 2J, K**), indicating that the effect was indeed due to the loss of *Ets21C* function. Thus, *Ets21C* is required for effective regeneration.

*Ets21C* is part of the Ets-family of DNA binding transcription factors that are broadly conserved in animals. The *Ets21C* mammalian orthologs are *Ets-related gene (ERG)* and *Friend Leukemia Integration 1 Transcription Factor (FLI1)*, both of which can act as proto-oncogenes (Kar and Gutierrez-Hartmann, 2013). In *Drosophila*, although *Ets21C* is not expressed in undamaged third instar wing discs, its expression is upregulated in tumorous imaginal discs (Kulshammer *et al*., 2015; Toggweiler *et al*., 2016) and it has been shown to be involved in adult midgut homeostasis (Jin *et al*., 2015; Mundorf *et al*., 2019). *Ets21C* is a downstream target of JNK/AP1 signaling in these contexts (Kulshammer *et al*., 2015; Toggweiler *et al*., 2016; Mundorf *et al*., 2019). Similarly, we found that even in undamaged discs, activation of the JNK pathway induces *Ets21C* expression (**supplemental fig. 8**). The JNK/AP1 pathway is known to be critical for regeneration (Bosch *et al*., 2005; Mattila *et al*., 2005; Katsuyama *et al*., 2015; Harris *et al*., 2016; Harris *et al*., 2020). Thus, we hypothesized that Ets21C could be functioning downstream of JNK/AP1 signaling to activate a regeneration-specific GRN in the blastema.

We investigated if *Ets21C* mutant tissues would fail to upregulate secreted molecules during regeneration. *Pvf1* and *upd3* are expressed in the inner region of the regenerative secretory zone. In wild-type regenerating discs, *Pvf1* and *upd3-lacZ* (Bunker *et al*., 2015) expression was detected within the center of the blastema and also within cellular debris (**Figure 3A, C**). In contrast, regenerating *Ets21C^-/-^* mutant discs showed a substantial decrease both in Pvf1 and *upd3-lacZ* expression (**Figure 3B, D**), indicating that *Ets21C* is required to initiate expression of both ligands in response to damage. In addition, we found that this reduced expression of *upd3* and possibly its paralogs impacted downstream JAK/STAT signaling. In control tissues, the STAT activity reporter was expressed in the center of the blastema (**Figure 3E**) as well as in the surrounding hinge regions where the JAK/STAT pathway is active during development (Ayala-Camargo *et al*., 2013; La Fortezza *et al*., 2016). In contrast, *Ets21C^-/-^* mutant tissues failed to activate the JAK/STAT reporter within the cells at the center of the blastema (**Figure 3F**), while JAK/STAT signaling in the hinge remained unaffected. Thus, *Ets21C* is required for localized expression of Pvf1 and Upd ligands within the inner regenerative secretory zone. The disruption of the inner regenerative secretory zone is also observed when we assess the pattern of cell proliferation in regenerating *Ets21C* mutant discs, which showed a reduced central non-proliferating zone (**Figure 3G, H; supplemental fig. 9**). This observation suggests that Ets21C is required for proper function of the inner regenerative secretory zone and potentially for its initial establishment.

**Figure 3.**
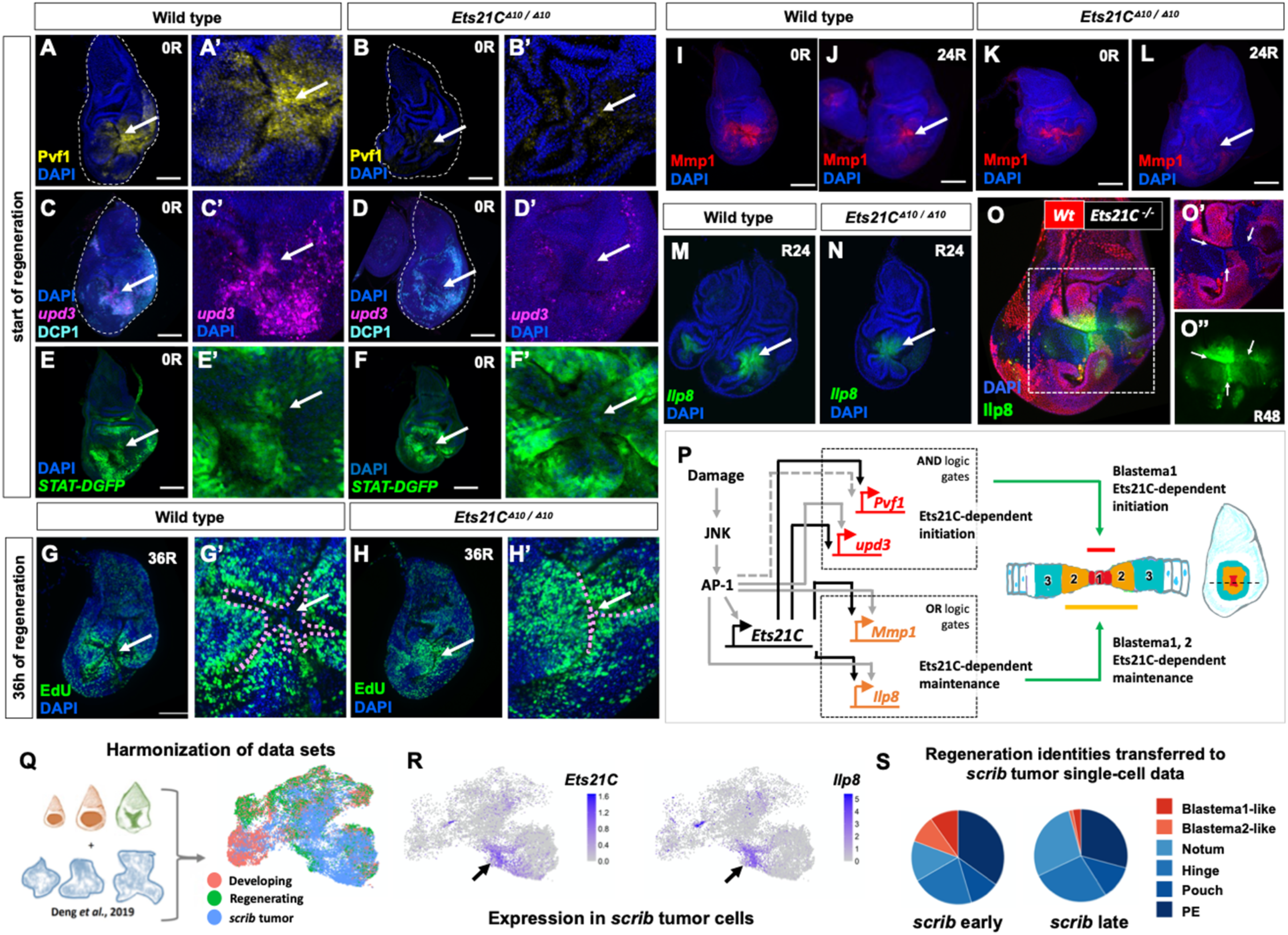
*Ets21C* is required to initiate and sustain a pro-regenerative program. (**A-N**) Wild-type or homozygous *Ets21C* mutant regenerating imaginal discs. (**A-F**) Imaginal discs at the start of the regeneration period (0R), (**A, B**) showing Pvf1 expression, (**C, D**) *upd3-lacZ* expression and apoptosis (cleaved Dcp-1), and (**G, H**) JAK/STAT activity (*STAT-DGFP* reporter). Note the decreased activation in *Ets21C* mutant tissues. (**G, H**) Cell proliferation as assessed by EdU incorporation at 36h of regeneration (see also **fig S9**). (**I-L**) Mmp1 expression at the start (0R) and after 24h of regeneration (24R). (**M, N**) *Ilp8-GFP* reporter expression at 24R. (**O**) Mosaic tissues created by mitotic recombination. *Ets21C* mutant cells marked by the absence of RFP. Note that *Ets21C* mutant cells are able to proliferate during regeneration, but show a cell-autonomous decrease in *Ilp8-GFP* expression at 48h of regeneration. (**P**) Model of GRN initiated and sustained by Ets21C during regeneration. (**Q-S**) Single-cell comparison of regenerating and *scrib* tumorigenic discs. (**Q**) Diagram of combined single-cell data. (**R**) Expression of *Ets21C* and *Ilp8* in *scrib* tumor cells. (**S**) Summary of cell types predicted in the *scrib* tumor cell dataset based on identity transfer from regeneration single-cell data. Note presence of Blastema-like cells within *scrib* tumors.

We next examined if Ets21C regulated the expression of genes expressed in the outer regenerative secretory zone. Wg expression during regeneration was unaffected in *Ets21C* mutants (**supplemental fig. 10**). While Mmp1 expression was observed at early stages of regeneration, it was prematurely absent by 24 hours in *Ets21C* mutants (**Figure 3I-L**), demonstrating that Ets21C is required to maintain Mmp1 expression during regeneration. Mmp1 is important for proper blastema formation (McClure *et al*., 2008) and effective regeneration (Harris *et al*., 2020). Similarly, *Ilp8* expression appeared normal at earlier time points of regeneration, but showed a slight decrease as regeneration progressed (**Figure 3M, N**). Regenerating discs, when compared to normal development, have a relatively immature transcriptional state (**supplemental fig. 11**), marked by the delay in the expression of the transcription factor Broad (Narbonne-Reveau and Maurange, 2019) (**supplemental fig. 11**). *Ets21C^-/-^* mutants, however, showed premature expression of Broad during regeneration (**supplemental fig. 11**). In addition, a regeneration-induced developmental delay that occurs in more distant tissues, specifically the eye disc, was also reduced in *Ets21C^-/-^*larvae (**supplemental fig. 11**). *Ilp8* is crucial for delaying pupariation (Colombani *et al*., 2012; Garelli *et al*., 2012), and this delay is correlated with regeneration outcomes (Smith-Bolton *et al*., 2009; Halme *et al*., 2010; Katsuyama *et al*., 2015; Harris *et al*., 2016). Indeed, *Ets21C* animals ended the larval phase of development approximately 30h before regenerating controls (**supplemental fig. 12**), which is likely the result of a decrease in Ilp8 levels and other signaling molecules. Thus, *Ets21C* mutants have both local and systemic defects in their regenerative response that collectively contribute to the reduced regeneration.

To test whether *Ets21C* regulates *Ilp8* cell-autonomously, we generated mosaic discs containing patches of both *Ets21C^-/-^* mutant and wild-type cells prior to tissue damage (see Materials and Methods). In regenerating discs, clones of *Ets21C* cells (marked by the absence of RFP) are comparable in size to wild-type clones, indicating that *Ets21C* does not have a cell-autonomous function in regulating cell proliferation during regeneration (**Figure 3O**). However, the *Ets21C^-/-^* mutant cells showed a cell-autonomous decrease in *Ilp8-GFP* expression after 48h of regeneration compared to wild-type cells (**Figure 3O’’**), indicating that as with *Mmp1*, Ets21C is required to sustain *Ilp8* expression. We propose that Ets21C works downstream of AP-1 in a type-1 coherent feed-forward loop (Alon, 2007), where the target genes that require Ets21C for initiating expression have enhancers with AND logic gates and target genes that require Ets21C for sustained expression during regeneration have enhancers with OR logic gates (**Figure 3P**). This GRN model suggests that *Ets21C* is critical for cells to interpret sustained JNK/AP-1 signaling and activate pro-regenerative pathways.

While *Ets21C* function is required for regeneration and not necessary for normal development, it is known to be expressed in tumorous imaginal discs that have mutations that disrupt apicobasal polarity (Kulshammer *et al*., 2015; Toggweiler *et al*., 2016). Moreover, in one study, reducing *Ets21C* function was shown to reduce overall tumor size (Toggweiler *et al*., 2016). Recent single-cell studies (Deng *et al*., 2019)(Ji *et al*., 2019) of tumorous imaginal discs caused by mutations in the apicobasal polarity regulator *scribble (scrib)* (Bilder *et al*., 2000) have demonstrated considerable cellular heterogeneity. By harmonizing our data with published single-cell RNAseq data derived from tumorous *scrib* discs (Deng *et al*., 2019) (**Figure 3Q; supplemental fig. 13**), we found that a subset of cell clusters have similar transcriptomes to the regenerative secretory zone (Blastema1 or Blastema2 cell clusters) (**Figure 3S**). Notably, these cell clusters express *Ets21C* along with *upd3, Pvf1, Mmp1*, and *Ilp8* (**Figure 3R; supplemental fig. 13**) and the cells are more prevalent at earlier stages of disc overgrowth (**Figure 3S; supplemental fig. 13**). Thus, while most cells in the disc have defects in apicobasal polarity, only a small subset of cells appear to activate this pro-regenerative GRN featuring *Ets21C*. Since reducing *Ets21C* function reduces growth in similar tumor models (Toggweiler *et al*., 2016), the presence of blastema-like cells could be critical for promoting the overgrowth of tumorous discs.

In conclusion, we have discovered a GRN that is dispensable for normal development yet essential for regeneration. Regeneration-specific GRNs may also exist in vertebrates and their reactivation could be valuable for regenerative medicine. Finally, the role of pro-regenerative GRNs in oncogenesis merits further exploration.

## Materials and Methods

### Single-cell data collection

For each sample, approximately 300 regenerating wing-imaginal discs were collected after 24 hours of regeneration. Regenerating discs were dissected within 1 hour in Supplemented Schneider’s Medium. The samples were then processed according to the protocol outlined in (Everetts *et al*., 2021). Briefly, we used a mixture of trypsin and collagenase to enzymatically dissociate the tissues. Then we used FACS to eliminate both apoptotic cells and cellular debris. Because our tissue dissociation protocol enriched for myoblasts, we decided to specifically sort out myoblasts during the collection of our second regeneration sample. This was done with a *Holes in muscle (Him)-GFP* construct that specifically labeled the myoblasts (Rebeiz *et al*., 2002). The myoblasts represented 75.67 % of the cells from the sample without *Him-GFP* and 12.71 % cells of the sample with *Him-GFP* (as determined by single-cell analysis). Single-cell suspensions were barcoded for single-cell RNA sequencing with the 10X Chromium Single Cell platform (v2 chemistry). Barcoded samples were sequenced on an Illumina NovaSeq (S2 flow cell) to over 60% saturation.

### Single-cell data analysis

Single-cell sequencing reads were aligned with the 10x Genomics CellRanger pipeline (v.2.2.0) to the *Drosophila melanogaster* transcriptome (version 6.24, FlyBase). Analysis of the single-cell data was conducted in the R and Python programming languages, primarily using the packages scvi-tools v0.9.1 (Gayoso *et al*., 2021) and Seurat v3 (Stuart *et al*., 2019).

Before cell filtering, we used scvi-tools to harmonize our single-cell data from regenerating wing discs with the single-cell data from developing wild-type wing discs presented in our previous study (accession number GSE155543) (Everetts *et al*., 2021). We used Seurat’s variance-stabilizing transformation method to select 1000 variable genes for each batch, and the scVI VAE model was trained on the union of these genes with the following parameters: n_latent = 15, n_layers = 2, gene_likelihood = “nb”, max_epochs = 400, and train_size = 0.8. The scVI latent space was used as the input for Seurat’s clustering algorithm (parameters: clustering k.param = 35, clustering resolution = 2.0; default parameters otherwise). Known transcriptional markers were used to classify cell clusters: *SPARC* and *twist* for AMPs, *Fasciclin 3(Fas3)* and *narrow* for the disc epithelium, and *regucalcin* and *Hemese (He)* for hemocytes (**supplemental fig. 2**). We also identified an unusual cell cluster that expressed both AMP and epithelium markers, with slightly elevated average nGene and nUMI counts, which suggested that these cells were actually doublets. When we applied the tool DoubletFinder (McGinnis *et al*., 2019) (parameters: 30 PCs, pN = 0.25, expected doublets (nExp) = 7.5% of total cells in each batch, and pK determined by the recommended BCmvn method) to each individual batch, the majority of cells within this cluster had been classified as potential doublets within their respective batches. We determined this cluster to represent AMP-epithelium doublets and removed it from subsequent analysis. We isolated the remaining disc epithelium and AMP clusters and filtered these cell types separately.

When filtering the disc epithelium cells, we first processed each batch using the standard Seurat pipeline (parameters: nfeatures = 1000, npcs = 30, k.param = 20, clustering resolution = 2.0; default parameters otherwise) and removed low-quality clusters. We classified low-quality clusters as having: 1) an average nGene less than 1 standard deviation below the average nGene of all cells, 2) an average percent.mito greater than 1 standard deviation above the average percent.mito of all cells, and 3) an abundance of negative marker genes compared to positive markers genes (as calculated by a Wilcoxon test). After removing low-quality clusters, we marked potential doublets within the epithelium cells by applying DoubletFinder (same parameters as described initially) to the epithelium cells in each batch. Batches were harmonized with scVI (same parameters as described initially), trained on the union of the top 1000 variable genes within the epithelium cells for each batch as determined by Seurat. After harmonization, we used the scVI latent space as a basis for Seurat clustering (parameters: k.param = 35, resolution = 2.0; default parameters otherwise). We removed a cluster that we determined to be epithelium-epithelium doublets, based on the following characteristics: (1) a noticeably higher average nGene compared to all other clusters (the only cluster with an average nGene > 1 standard deviation above the average nGene of all cells), (2) an extreme abundance of potential doublets as classified by DoubletFinder from each batch (~70% of all potential doublets classified were contained within this cluster), and (3) a lack of marker genes (both positive and negative) when compared to other clusters. We also removed a cluster that we determined to represent a small number of trachea cells, based on the unique expression of marker genes tracheal-prostasin and waterproof. We re-ran our variable gene selection, scVI harmonization, and Seurat clustering. Data was visualized in 2 dimensions with UMAP (parameters: min.dist = 0.1; default parameters otherwise).

### Gene signature analysis of the blastema

For each identity combination (hinge-pouch, pouch-notum, and notum-hinge), gene signatures were constructed as follows: First, differential expression was performed between wild-type (non-regenerating) cells of each identity pair (e.g., for the hinge-pouch signature, differential expression was performed between cells from (Everetts *et al*., 2021) classified as hinge vs. cells classified as pouch). This was conducted using a Wilcoxon test via Seurat’s FindMarkers function, selecting genes with a natural-log fold-change of greater than 0.25 (logfc.threshold = 0.25) and a Bonferroni-corrected p-value of < 0.05. This provided three gene sets that differentiated hinge-pouch, pouch-notum, and notum-hinge identities. Second, principal component analysis was performed on all cells using each gene set. The first principal components from each analysis were defined as the gene signatures, as they best separated cells of the different identities. The signature scores of cell clusters were visualized in 2-dimensions using Seurat’s VlnPlot function (**Figure 1J; supplemental fig. 5A, B**) and in 3-dimensions using the R package Plotly (**supplemental fig. 5C**).

### Gene signature of cellular maturity

To determine the relative cellular maturity (or developmental progression) of individual cells within the regenerating tissue we generate a gene signature score based on genes with differential expression during normal development. First, we selected genes with consistent differential expression between epithelial cells from mid (younger) and late (older) 3rd instar imaginal discs (with a threshold of greater than 0.25 natural-log fold-change). Second, this gene set was then used to perform principal component analysis to derive a cellular maturity score. The relative cellular maturity score of cells from normal developing and regenerating discs were visualized on the UMAP (**supplemental fig 11B**).

### Single-cell comparison of regenerating and scrib tissues

The expression matrices for the *scrib* single-cell data were downloaded from GEO, accession number GSE130566 (Deng *et al*., 2019). Gene names were updated to match those within our regeneration and wild-type datasets. All *scrib* datasets (4d, 5d, 8d, and 14d) were harmonized with scVI (n_latent = 15, n_layers = 2, gene_likelihood = “nb”, max_epochs = 400, and train_size = 0.8), trained on the union of the top 1000 variable genes for each batch as determined by Seurat. Clustering was performed using Seurat, and we isolated the *scrib* epithelium clusters (identifiable by high expression of *Fasciclin 3* and *narrow*) for subsequent comparison with the regeneration and wild-type epithelium data. No *scrib* epithelium cells were filtered during this comparative analysis.

The epithelium data from regeneration, wild-type, and *scrib* samples was initially harmonized with scVI (n_latent = 15, n_layers = 2, gene_likelihood = “nb”, max_epochs = 400, and train_size = 0.8), trained on the union of the top 1000 variable genes for each batch as determined by Seurat. The weights from this scVI model were used to initialize a scANVI model (using the from_scvi_model function) for semi-supervised training and label transfer. The cluster identities from our regeneration analysis (**Figure 1B**) were supplied as input labels (via setup_anndata), with all *scrib* cells marked as “Unknown”. The scANVI model was trained for 50 epochs (max_epochs = 50) to predict the probability of “Unknown” cells belonging to each of the input labels. Input labels were subsampled during training (n_samples_per_label = 150) to prevent the loss of labels with relatively few cells (e.g., Blastema1) during label transfer. After training, the scANVI latent space was used as a basis for UMAP (**supplemental fig. 13B**), and the transferred labels corresponded to the highest predicted identity for each cell (**supplemental fig. 13C**).

### Immunohistochemistry and microscopy

The following antibodies were from the Developmental Studies Hybridoma Bank (DSHB): mouse anti-Wg (1:100, 4D4), mouse anti-Mmp1 (1:100, a combination of 14A3D2, 3A6B4 and 5H7B11), mouse anti-Broad-Z1 (BrZ1) (1:100, Z1.3C11.OA1), and rat anti-Elav (1:50, Elav-7E8A10). The following antibodies were gifts: rat anti-Zfh2 (1:100, Chris Doe (Tran *et al*., 2010)), rat anti-Twist (1:1000, Eric Wieschaus), rat anti-Pvf1 (1:500, Ben-Zion Shilo (Rosin *et al*., 2004)), and pan-hemocyte anti-H2 (1:100) (Kurucz *et al*., 2003). The following antibodies are from commercial sources: rabbit anti-cleaved Death caspase-1 (Dcp-1) (1:250, Cell Signaling); chicken anti-GFP (1:500, ab13970 Abcam, Cambridge, UK); rabbit anti-PHH3 (1:500, Millipore-Sigma). Secondary antibodies were from Cell Signaling. Nuclear staining with DAPI (1:1000). Tissues were imaged on a Zeiss Axioplan microscope with Apotome attachment, using 10x and 20x objectives. Image files were processed with ImageJ software.

### EdU assay and quantification

For EdU staining, live discs were incubated in Schneider’s medium (ThermoFisher 21720024) with EdU for 30 minutes, following the protocol for the Click-iT EdU Cell Proliferation Kit, Alexa Fluor 555 (ThermoFisher C10338) and Alexa Fluor 647 (ThermoFisher C10340). After the incubation, discs were fixed in 4% PFA for 15 min, before proceeding with standard antibody staining, as detailed above.

EdU intensity was quantified using the ImageJ software. For each regenerating disc, a square box was drawn, centered around the blastema. The length of the box was 140 microns for the 0h R discs, 160 microns for the 24h and 36h R discs, and 200 microns for the 48h R discs. The EdU intensity was measured at every pixel along the two diagonals of each box using ImageJ’s “Plot Profile” function. Subsequent analysis was done using R software. The measured EdU intensities were first z-normalized (i.e., for all values in a measured profile, subtract the mean and divide by the standard deviation) and then averaged across all diagonals from all processed discs at each regenerating time point. The average normalized (scaled) EdU intensity was plotted with the package ggplot2, and smoothed curves were added using the stat_smooth function with method = “gam”.

### Drosophila stocks and husbandry

The stocks that were used in this study include: *Ets21C^Δ10^* (Mundorf *et al*., 2019); *upd3-lacZ* (Bunker *et al*., 2015); *eyFLP; arm-lacZ FRT40A; hsFLP; FRT40A; hsFLP; FRT40A ubi-RFP; rn-GAL4, tub-GAL80^ts^, UAS-rpr (Smith-Bolton *et al*., 2009); UAS-his2A::RFP;* and *Him-GFP* (Rebeiz *et al*., 2002). Stocks obtained from the Bloomington Stock Center include: *Ilp8-GFP (Ilp8^MI00727^*, Bl33079); *10XSTAT-DGFP (10XSTAT92E-DGFP*, Bl26199, Bl26200) (Bach *et al*., 2007); *Ets21C-GFP (Pbac-Ets21C-GFP.FLAG^VK00033^*, Bl38639)*; hh-Gal4; rn-Gal4* (Bl7405); *rn-GAL4, tub-GAL80^ts^, UAS-egr* (Bl51280)*(4)*; *Df(2L)BSC456* (Bl24960); *UAS-hepWt* (Bl9308)*; Ubi-FRT-stop-FRT-GFP^nls^* (BL32251) (Evans *et al*., 2009) *; lexAOp-FLP* (Bl55819); and *GMR26E03-lexA* (Bl54354) (Pfeiffer *et al*., 2010).

### Regeneration experiments

Unless otherwise noted, the genetic ablation system used to study regeneration was *rn-GAL4, tub-GAL80^ts^, UAS-eiger* (Smith-Bolton *et al*., 2009). Genetic ablation experiments were conducted by synchronizing development by collecting eggs on grape plates and picking 55 L1 larvae into vials with yeast paste. Temperature shifts to induce ablation (from 18°C to 30°C) were conducted on day 7 after egg lay (AEL) for 40 hours. The extent of adult wing regeneration was scored by binning the resulting wings into 5 categories (0%, 25%, 50%, 75%, and 100%) *(4)*. The resulting regeneration scores were calculated per population. Experimental replicates were done on separate days with a minimum of 2 vials per genotype and three replicates per genotype. Statistical comparison performed on regeneration scores using ANOVA followed by Tukey test for significance.

### Mitotic clones during regeneration

Mosaic tissues were generated by recombinase-driven (FLP/FRT) mitotic recombination within the genetic background of the ablation system. The expression of *hsFLP* was induced by an 1h heat-shock at 37°C on day 3 AEL, which generated clones throughout the imaginal discs prior to genetic ablation and regeneration. Mutant cells were labeled by the absence of RFP and wild-type cells were marked by 2X RFP. The genotype of the experimental larvae used to generate *Ets21C* mutant clones during regeneration: *hsFLP; Ets21C^Δ10^, FRT40A / ubi-RFPnls, FRT40A; rn-GAL4, tub-GAL80^ts^, UAS-eiger/Ilp8-GFP* (**Figure 3O**).

### Lineage-tracing experiments

We identify an enhancer for the gene *grain (grn)* that was primarily expressed in the inner-hinge, *GMR26E03-lexA* (Pfeiffer *et al*., 2010) during normal development (**Figure 1N**). Lineage-tracing was performed by permanently labelling the cells that expressed *GMR26E03-lexA* by driving the expression of the recombinase FLP *(lexAop-FLP)* to induce the removal of a stop-cassette *(Ubi-FRT-stop-FRT-GFP^nls^)* (**Figure 1M-O**).

### Pupariation timing experiments

Images were taken every 20 minutes of vials that contained animals as they transitioned between larva to pupa. This was performed at 18°C with a wide-angle camera (Arducam). Pupariation was scored by observing when the animals stopped moving and darken in color.

### Physical wounding assay

Wing discs were physically wounded *in situ* as described in (Yoo *et al*., 2016). Briefly, L3 larvae with the wing pouch fluorescently labeled (*rn-GAL4, UAS-his2A::RFP*) were visualized using a fluorescence microscope. The right wing pouch was wounded by carefully applying pressure on the larval cuticle using a thin gauge insulin needle without penetrating the larval cuticle. Larvae were then returned to vials containing Bloomington food and dissected 6 hours or 24 hours later.

## Acknowledgments

The authors would like to thank Mirka Uhlirova, Ben-Zion Shilo, David Bilder, Chris Doe, and Eric Wieschaus for stocks and reagents, current and former members of the Hariharan and Yosef labs for feedback, David Bilder and Craig Miller for helpful feedback on the manuscript, Shrey Saretha and Joel Sadler for technical assistance, and the Bloomington Stock Center, DRSC/TRiP Functional Genomics Resources, and Developmental Studies Hybridoma Bank for stocks and reagents.

## Funding

National Institutes of Health grant R35 GM122490 (IKH)

Chan-Zuckerberg Initiative 2018-184034 (NY)

## Competing interests

Authors declare that they have no competing interests.

## Data and materials availability

The single-cell RNA sequencing data presented in this study and analysis code will be made available upon publication.

## Supplementary Figures

**Supplementary figure 1.**
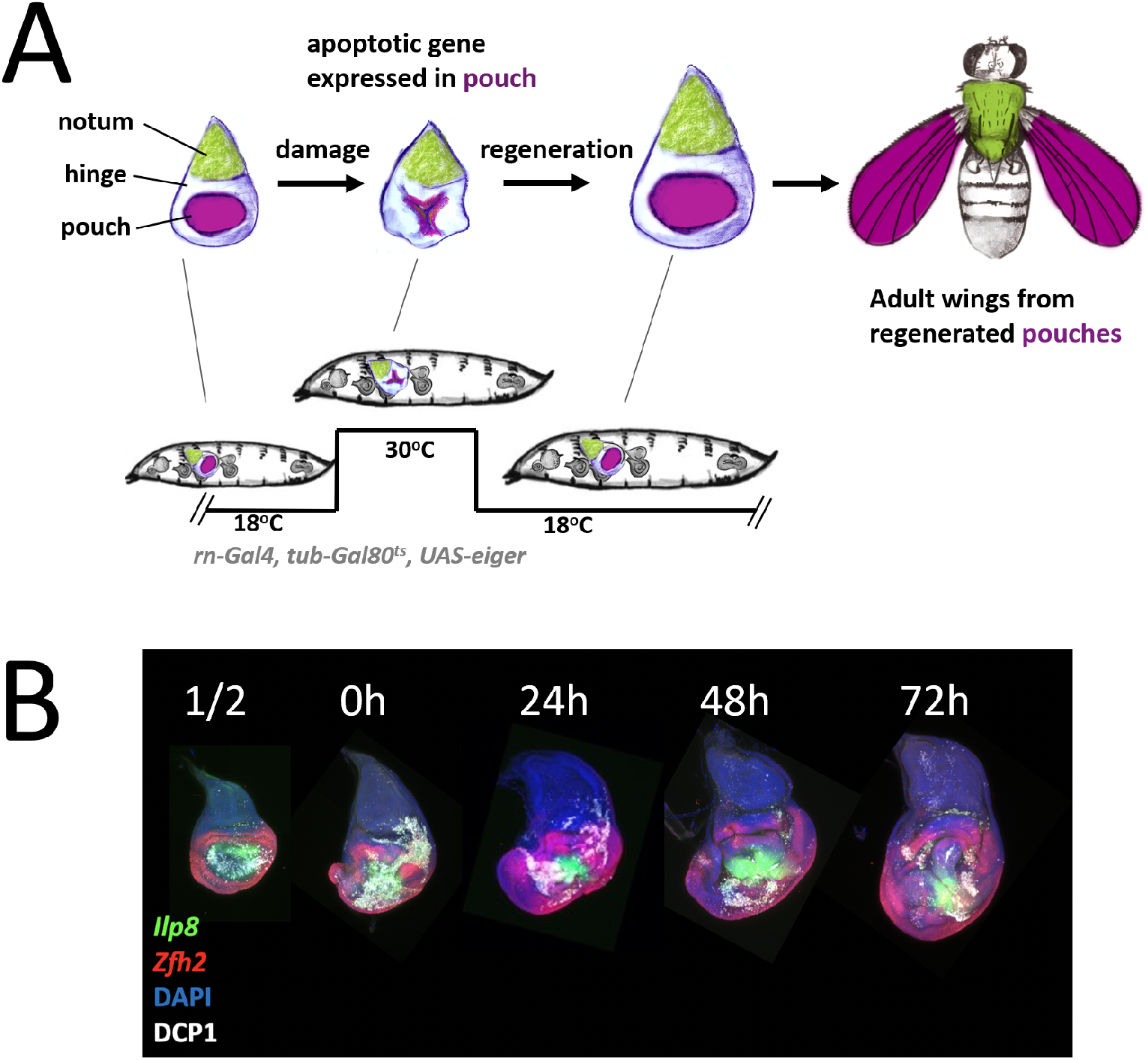
Schematic of genetic ablation system to study imaginal disc regeneration. (**A**) Schematic of genetic ablation system. The canonical domains of the wing disc and the adult structures to which they give rise are colored in green (notum), white (hinge), and purple (pouch). Expression of the pro-apoptotic gene *eiger* is targeted to the wing pouch using *rn-GAL4* and *UAS-eiger*. Gal4 function is inhibited at 18° C and permitted at 30°C by a ubiquitously-expressed temperature-sensitive Gal80 (*tub-Gal80^ts^*). (**B**) Imaginal discs can be dissected and analyzed during and after the ablation period. The half-way point through the 40h ablation is indicated by “½”. Other times refer to the time after the downshift to 18° C - the phase when regeneration occurs.

**Supplementary figure 2.**
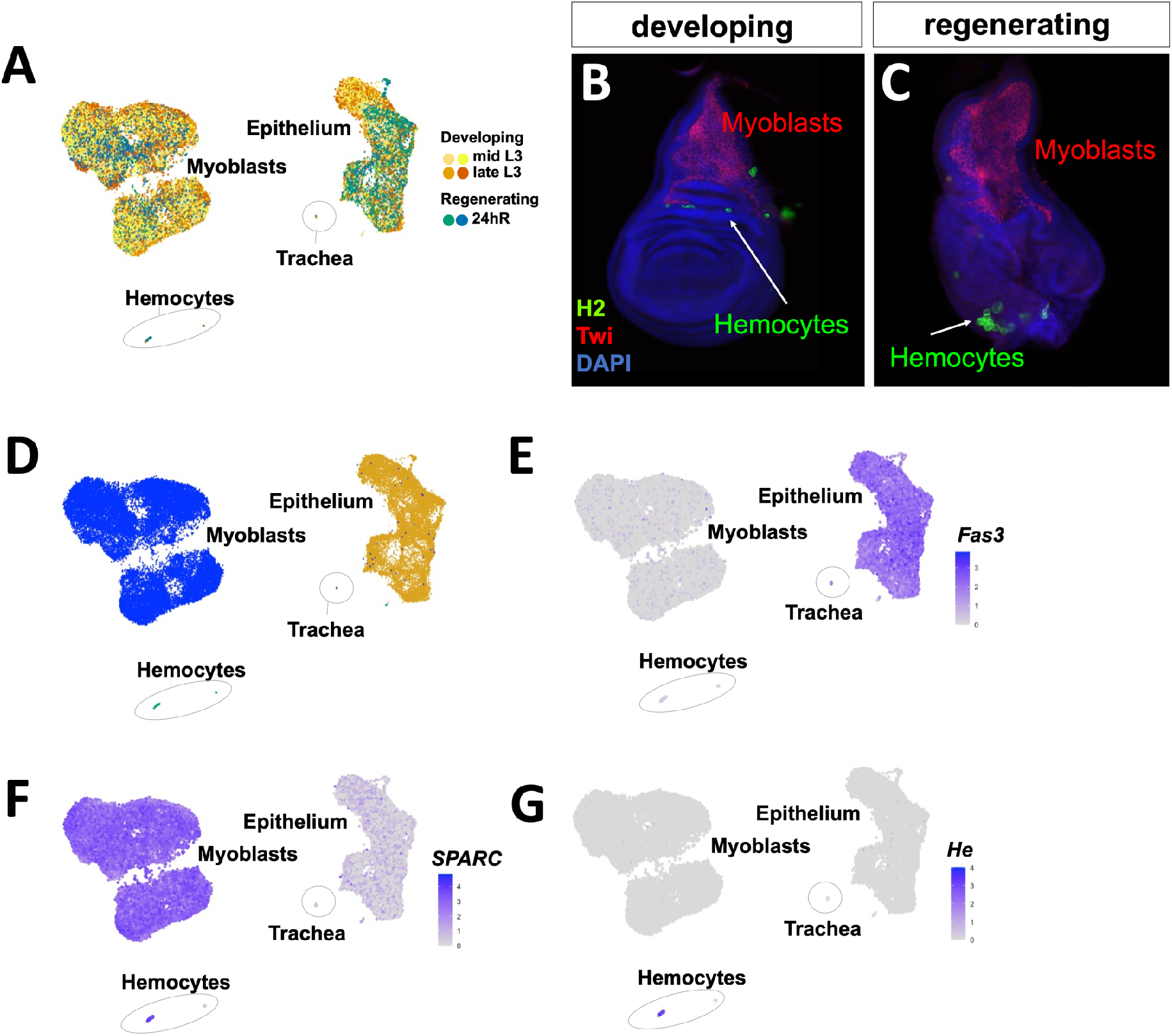
Three major cell types were identified from scRNAseq of wing imaginal discs. (**A**) Harmonized UMAP of scRNA data from wing imaginal discs. Data colored by sample of origin and labeled. Samples were derived from developing discs at the middle at late stages of the third larval instar (L3), as described previously (Everetts *et al*., 2021), and from regenerating discs 24h after the downshift to 18° C. Two biological replicates were obtained for each sample (see Materials and Methods). The three major cell types identified were epithelial cells, myoblasts and hemocytes. In addition, a few trachea cells were also identified. The cell counts from the regenerating discs were: 6,613 epithelial cells, 7,466 myoblasts, 224 hemocytes and 17 trachea cells. (**B, C**) Wing-imaginal discs stained with anti-H2 to label the hemocytes and anti-Twist to label the myoblasts in developing (**B**) and regenerating (**C**) wing discs. (**D**) UMAP colored by major cell types: myoblasts, epithelial cells, and hemocytes are shown in different colors (**E-G**) Expression of marker genes for the three major cell types: (**E**) *Fasciclin 3 (Fas3)* expression marks the epithelium; (**F**) *Secreted protein, acidic, cysteine-rich (SPARC)* expression marks the myoblasts; and (**G**) *Hemese (He)* marks the hemocytes.

**Supplementary figure 3.**
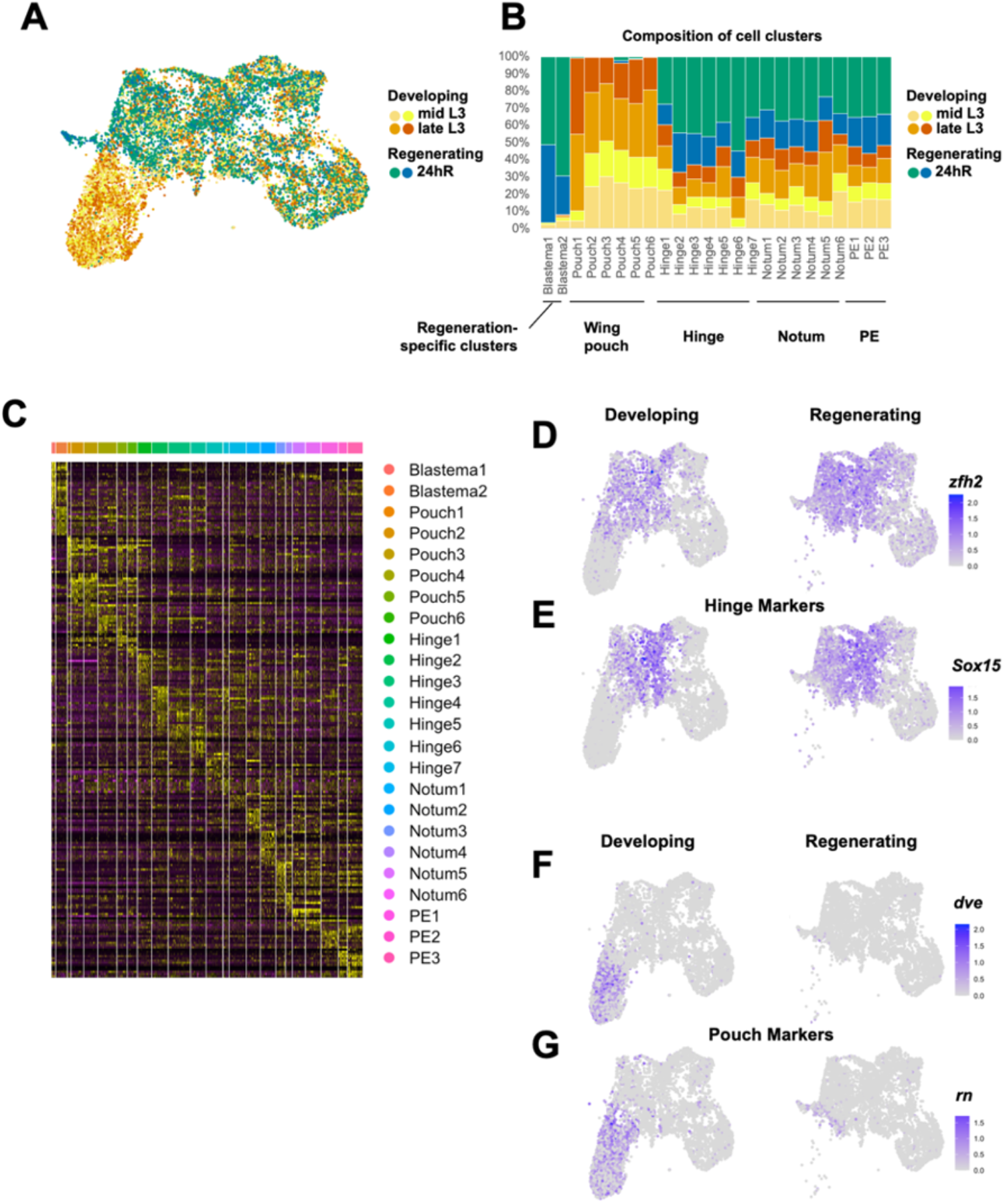
Composition of cell clusters in developing and regenerating discs. (**A**) UMAP of harmonized data from epithelial cells from regenerating and developing (from two time points) samples. Each dataset, including replicates, is represented in a distinct color. (**B**) Composition of cell clusters, as shown in **Figure 1B**. Note the underrepresentation of cells assigned to pouch clusters in regenerating discs and the near absence of cells assigned to the Blastema1 and Blastema2 clusters in developing discs. (**C**) Heatmap showing differential expression of marker genes through the different cell clusters of the harmonized epithelial cell object. Gene expression with individual cells from the single-cell data for the hinge makers *Zn finger homeodomain 2 (zfh2)* (**D**) and *Sox box protein 15 (Sox15)* (**E**) and for the pouch markers *defective proventriculus (dve)* (**F**) and *rotund (rn)* (**G**).

**Supplementary figure 4.**
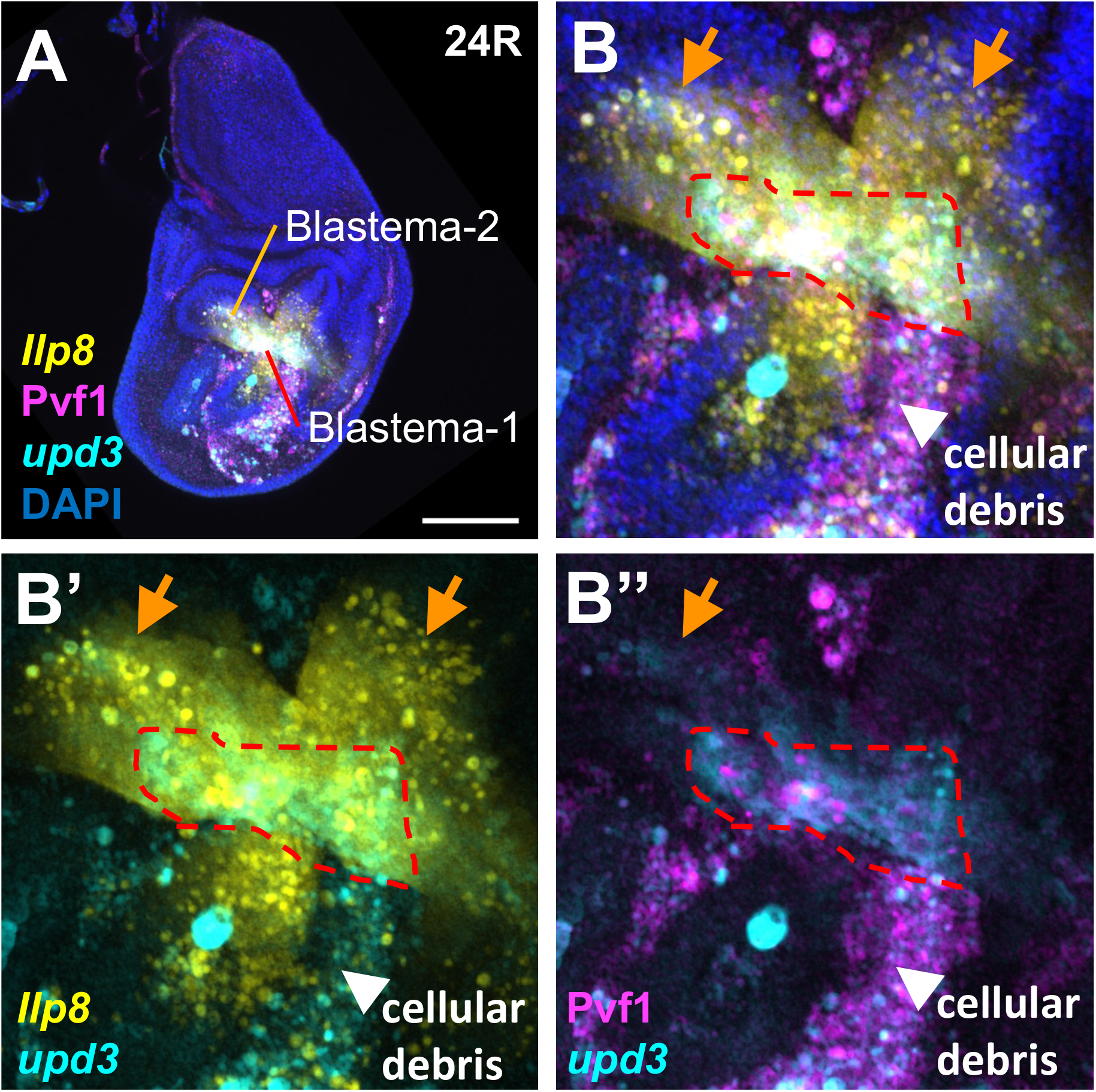
Co-expression of Blastema-1 and Blastema-2 marker genes within the regenerating epithelium. (**A, B**) Regenerating wing-imaginal disc, after 24h of regeneration, with transcriptional reporters and antibody staining to highlight the nested position of Blastema-1 and Blastema-2 cells within the regenerating epithelium. *Ilp8-GFP* expression is shown in yellow, anti-Pvf1 stain shown in magenta, and *upd3-lacZ (Bunker et al., 2015)* shown in cyan. (**B**) Magnification of blastema. Orange arrow highlights the region of *Ilp8* expression and the red dotted line highlights the region of higher Pvf1 and *upd3-lacZ* expression that is in the surviving epithelial cells. Note that the cellular debris shows evidence of expression of all three of these marker genes (white arrowhead). Microscopy scale bars = 100 μm.

**Supplementary figure 5.**
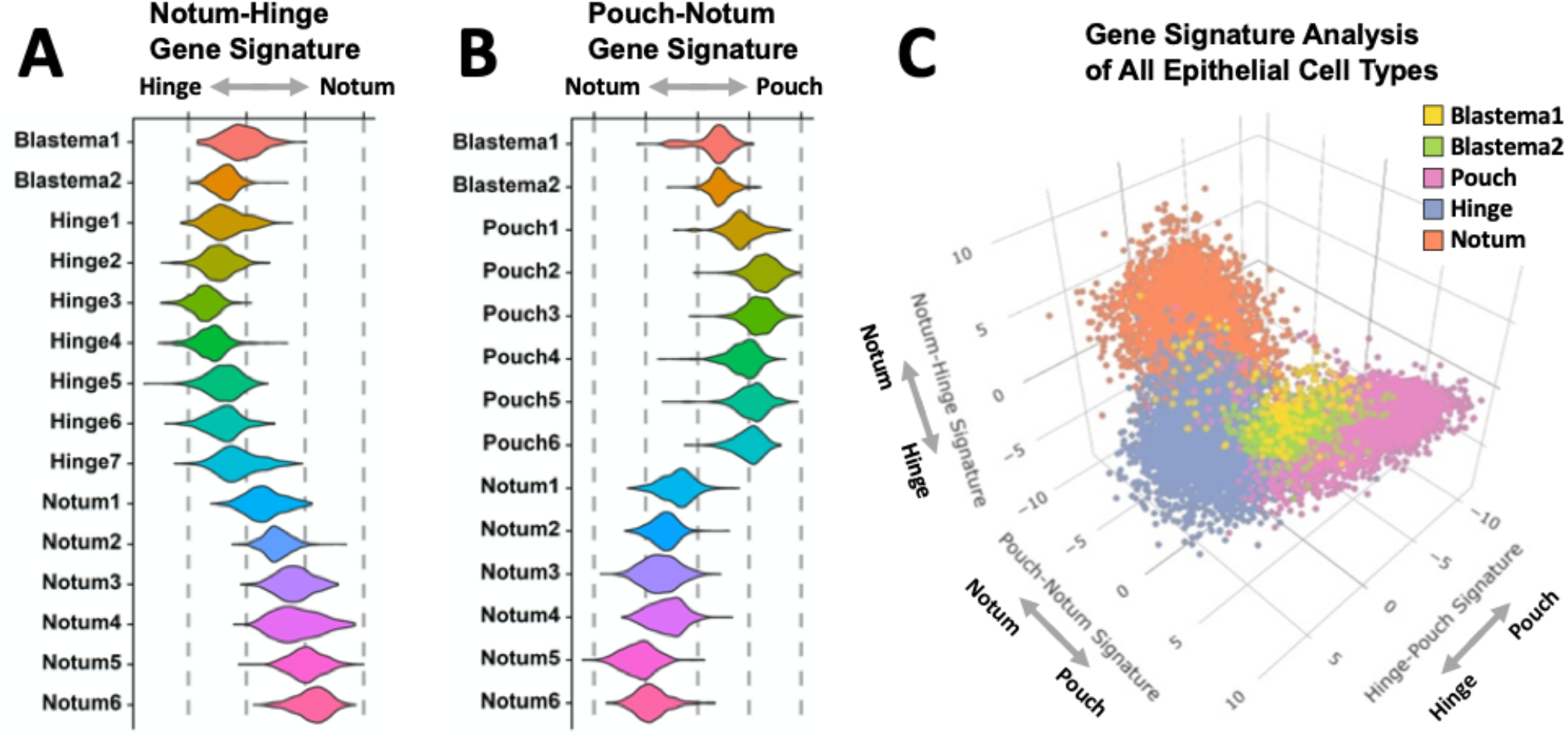
Comparative gene signature analysis of major epithelial domains for the blastema clusters. (**A, B**) Clusters within the scRNAseq dataset scored with a notum-hinge (**A**) or pouch-notum (**B**) gene signature. Note that scores for the blastema clusters are more closely aligned with the hinge (**A**) or pouch (**B**) rather than the notum. The hinge-pouch signature is shown in **Figure 1J.** (**C**) 3D signature plot of scRNAseq clusters scored by notum-hinge, pouch-notum, and pouch-hinge gene signatures. Axes correspond to values shown in **A**, **B**, and **Figure 1J**. Note that both Blastema1 and Blastema2 cells are centered between hinge and pouch fates.

**Supplementary figure 6.**
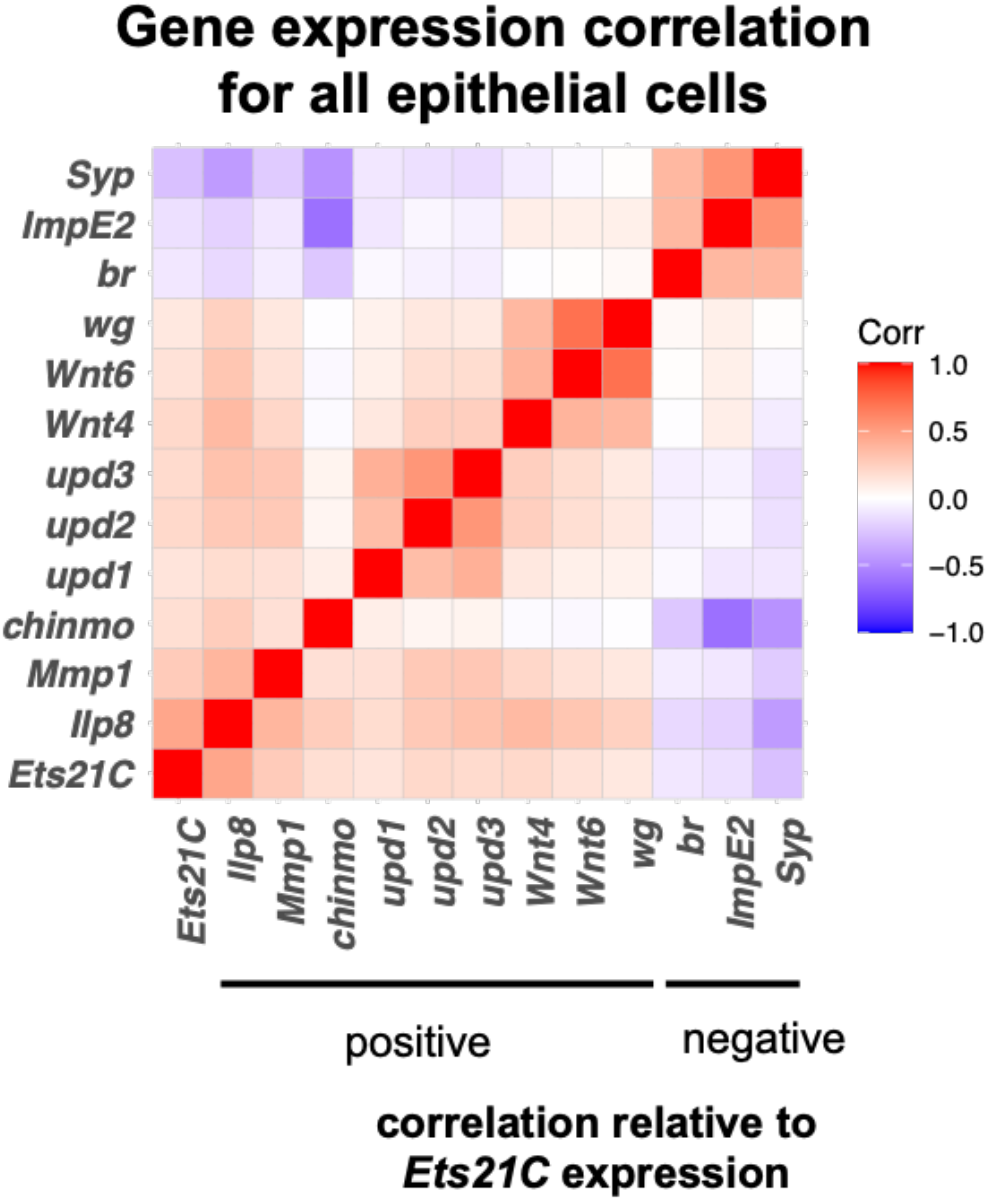
Genes that positively or negatively correlate with *Ets21C* expression. Pearson correlation of normalized gene expression data within all epithelial cells (both developing and regenerating datasets). The gene with the most correlated expression to *Ets21C* expression is *Ilp8. Ets21C* expression is also positively correlated with the expression of *Mmp1, chinmo, upd1, upd2, upd3, Wnt4, Wnt6* and *wg*. Examples of genes that show negative gene expression correlated with *Ets21C* include *broad (br), Ecdysone-inducible gene E2 (ImpE2)*, and *Syncrip (Syp)*.

**Supplementary figure 7.**
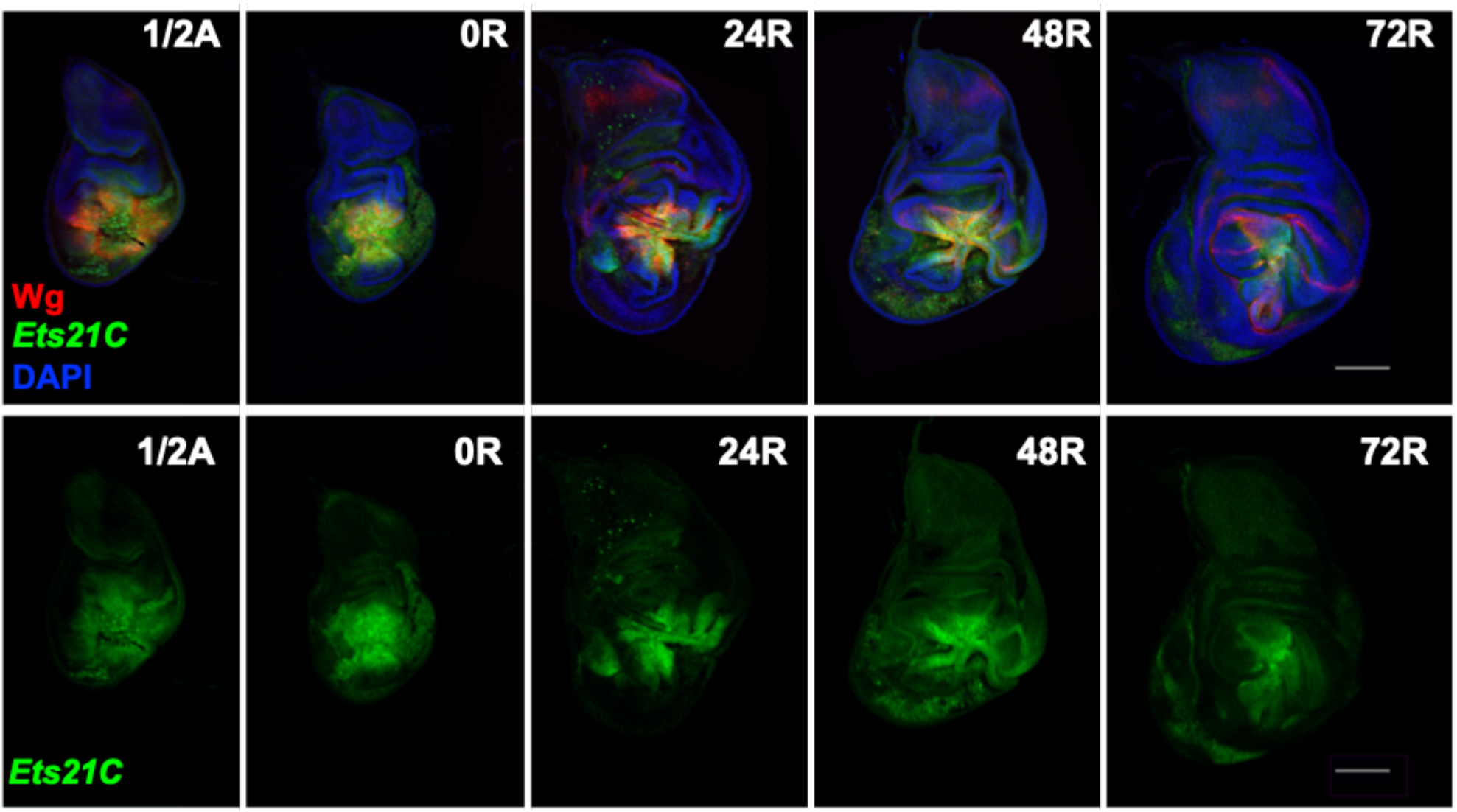
Expression of *Ets21C-GFP* over the course of regeneration. Wing imaginal discs dissected half-way (20 h) through the ablation period (1/2A) and time points (indicated in hours) during regeneration at 18C (0R, 24R, 48R, 72R). Regenerating wing discs are also stained with anti-Wg (in red). Regeneration is near complete by 72 h. Microscopy scale bars = 100 μm.

**Supplementary figure 8.**
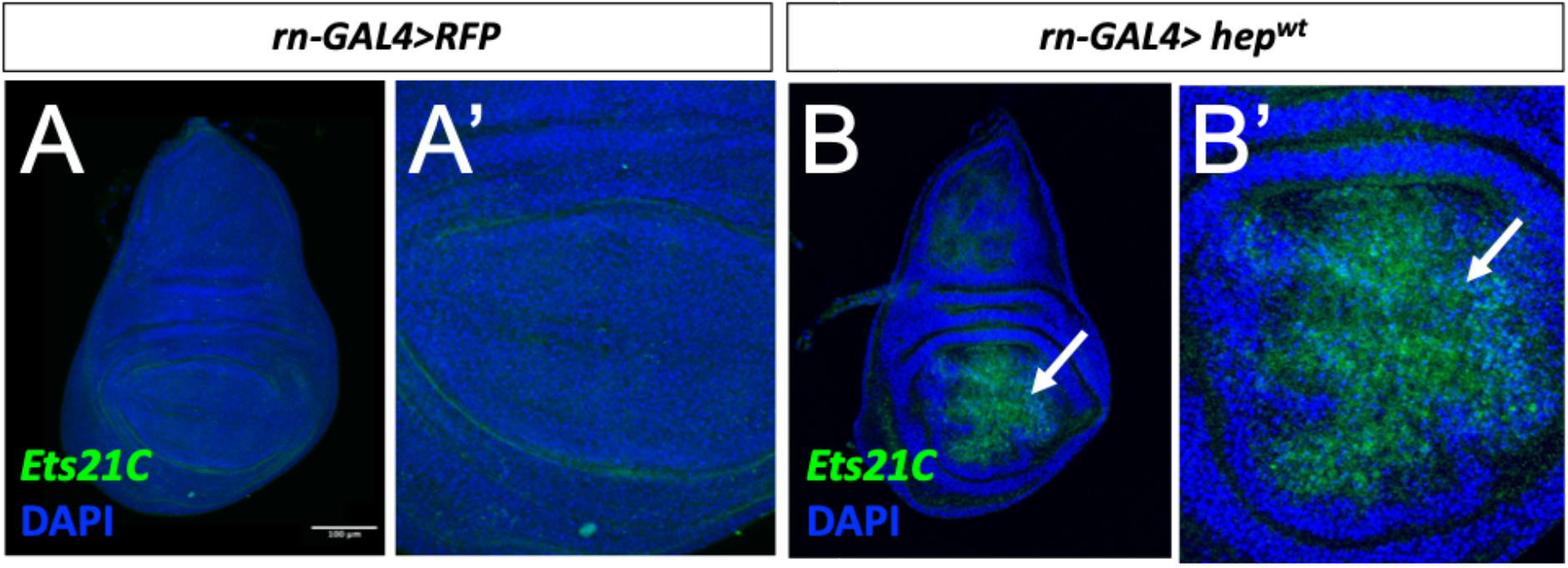
Activation of the JNK/AP1 pathway in non-ablated tissue induces the expression of *Ets21C*-GFP. (**A, B**) Expression of *Ets21C-GFP* in a wing disc with *rn-GAL4* driving the expression of *UAS-RFP* (**A**) or *UAS-hep^wt^* (**B**). Note that driving the expression of the wild-type version of the JNK-kinase *hemipterous (hep)* within the wing pouch leads to the expression of *Ets21C*. Microscopy scale bars = 100 μm.

**Supplementary figure 9.**
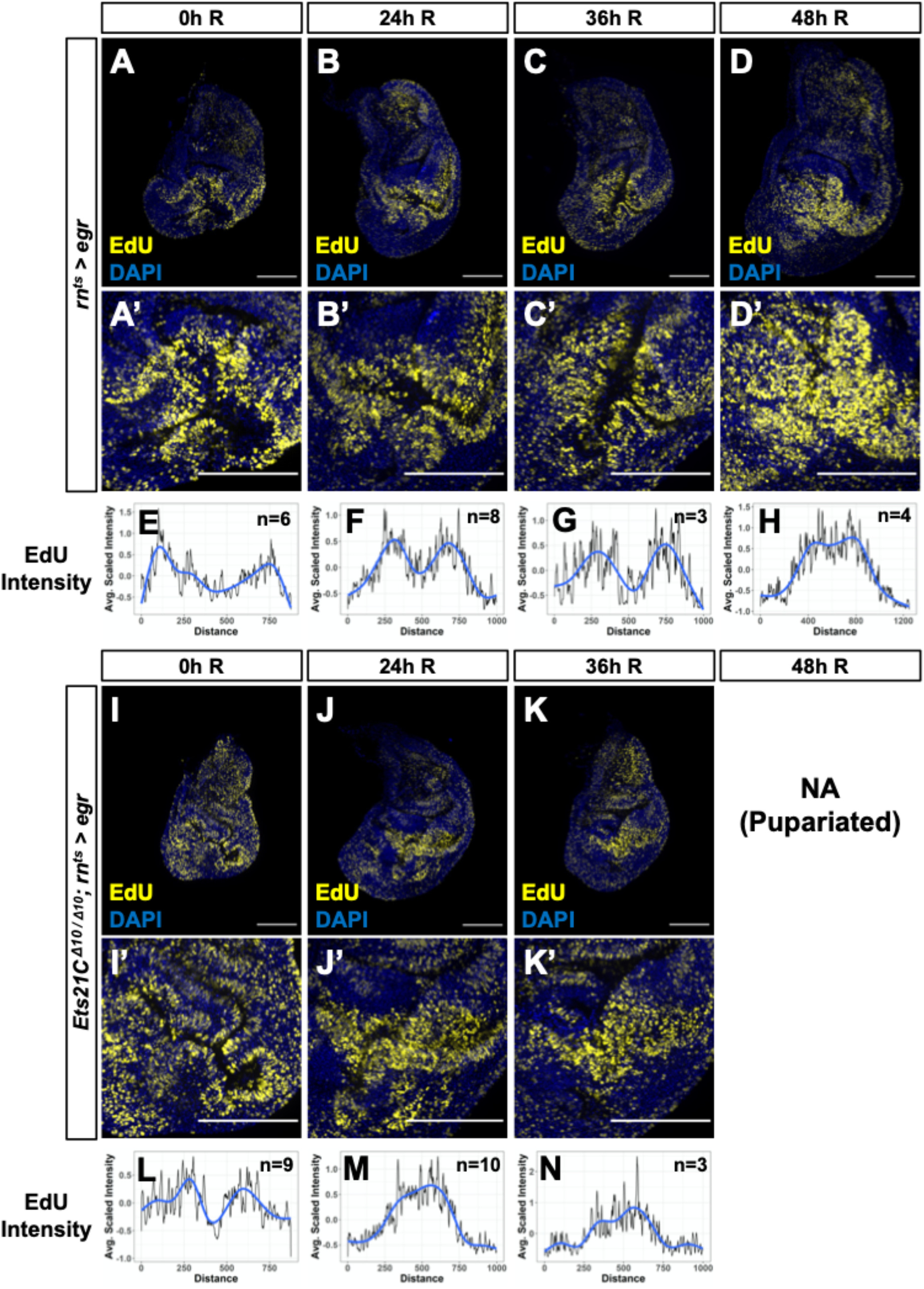
Cell proliferation during regeneration for wild type and *Ets21C* mutant wing discs. (**A-D, I-K**) Imaginal discs at different time points during regeneration (0R, 24R, 36R, 48R) for wild-type (**A-D**) or *Ets21C* mutant (**I-K**) tissues. Cell proliferation is marked by the incorporation of the thymidine analog EdU. Note that for the *Ets21C* mutants, the regeneration period is ended prematurely by pupariation (**Supplementary fig. 12**). (**E-H, L-N**) Profiles of average EdU intensity within the blastema at each regeneration time point for wildtype (**E-H**) or *Ets21C* mutant (**L-N**) discs. The y-axis corresponds to the average of z-normalized EdU intensity (see Materials and Methods). The x-axis corresponds to measured pixels centered around the blastema (i.e., the center of the distance axis is the center of the blastema). Note that in the wild-type tissues there is reduced EdU intensity within the center of the blastema, as evidenced by the bimodality of the EdU intensity profiles (most notable at 0R, 24R, and 36R). This pattern of proliferation is less robust within *Ets21C* mutant tissue, as evidenced by the unimodality of the EdU intensity profiles at 24R and 36R.

**Supplementary figure 10.**
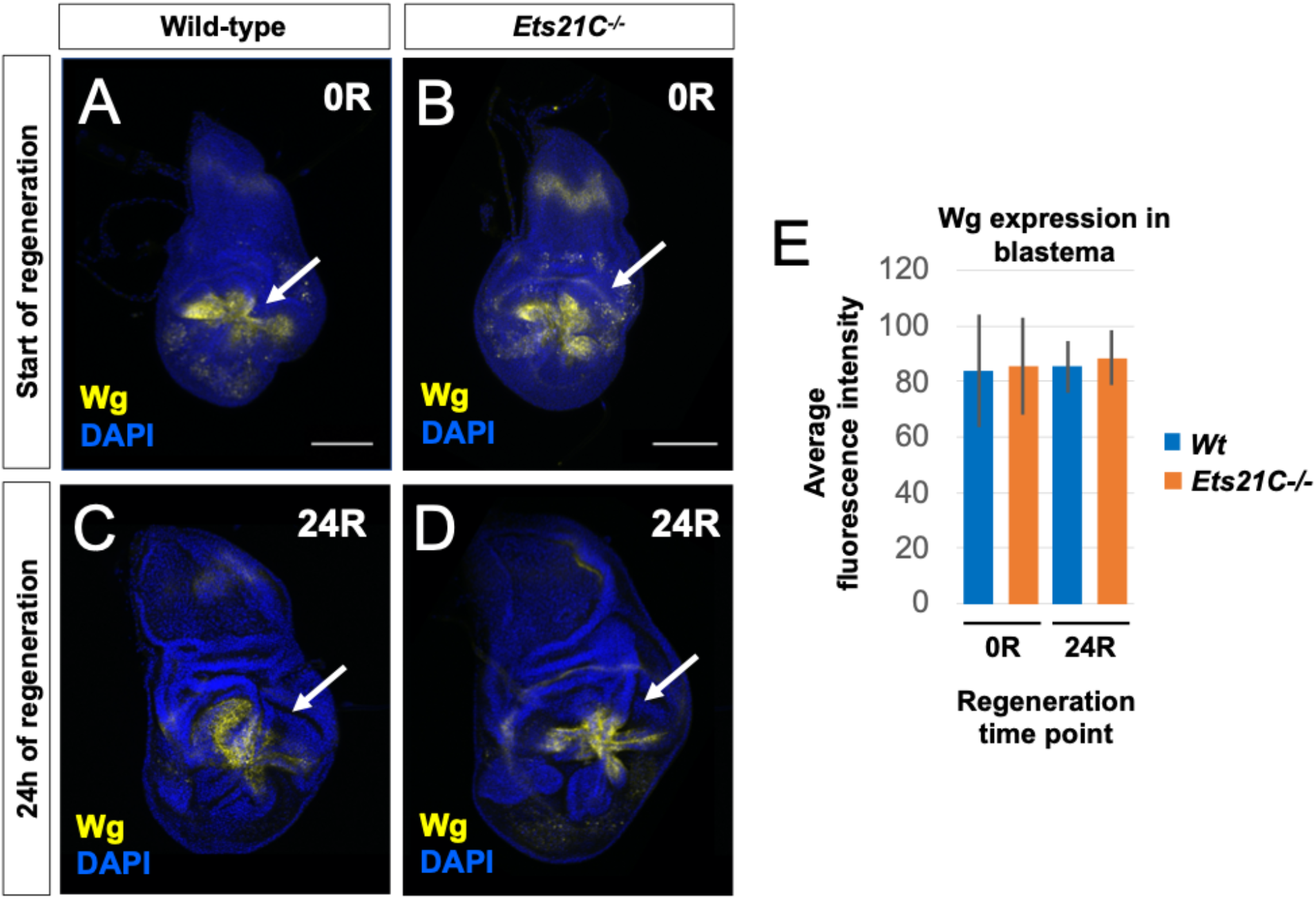
Wnt ligand Wg expression is unaffected during regeneration in *Ets21C* mutants. (**A-D**) Regenerating wing imaginal discs at the start (**A, B**) and after 24 h of regeneration (**C**, **D**) for wild type and *Ets21C* mutant tissues. Discs stained with anti-Wg and DAPI. Arrows are pointing to the area of the blastema. (**E**) Average fluorescent intensity of the anti-Wg stain within the blastema region for both wild type (Wt) and *Ets21C* mutant tissues. No change in the amount of Wg expression was detected at either regeneration time point.

**Supplementary figure 11.**
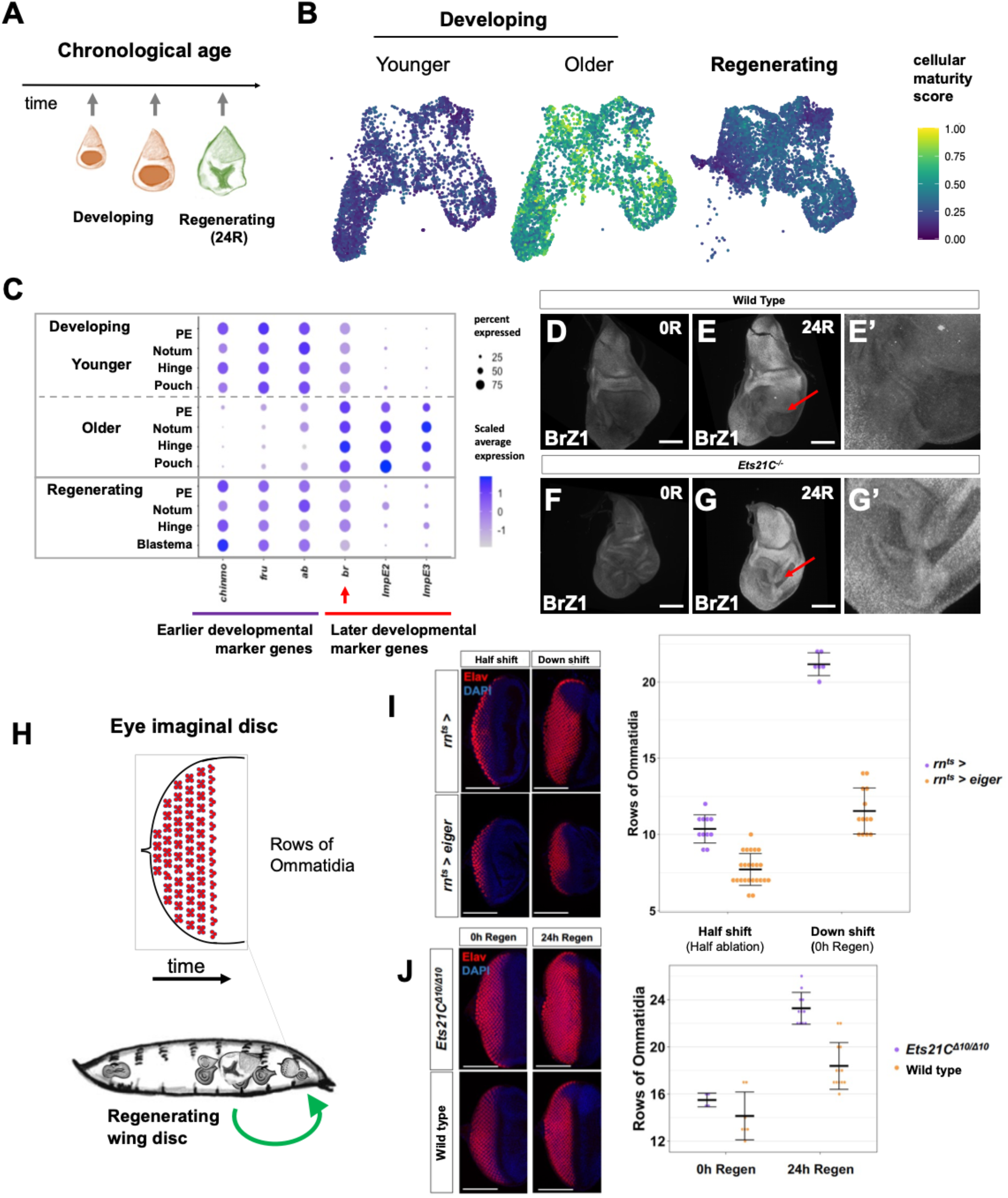
Developmental progression and cellular maturity during regeneration. (**A**) Diagram of the relative chronological age of our singlecell datasets. Note that the regenerating imaginal discs are chronologically older than both developmental time points, which are from mid and late 3rd instar, because tissue damage results in an extended larval phase during with regeneration occurs (Smith-Bolton *et al*., 2009; Halme *et al*., 2010; Katsuyama *et al*., 2015; Harris *et al*., 2016). (**B**) Based on genes with high differential expression during normal development, we calculated a cellular maturity score to determine the relative developmental maturity of individual cells from the single-cell data. Note that the cells within the regenerating sample show a lower or intermediate cellular maturity score, indicating that the developmental progression of the cells within the epithelium have been either paused or rejuvenated to an earlier time point in development. (**C**) Dot plot showing the relative expression levels for several genes that change over the course of development, including three genes expressed earlier development: *Chronologically inappropriate morphogenesis (chinmo), fruitless (fru)* and *abrupt (ab);* and three gene expressed in older imaginal discs during normal development: *broad (br), Ecdysone-inducible gene E2 (ImpE2)*, and *Ecdysone-inducible gene E3 (ImpE3)*. Note that the regenerating sample shows similar expression patterns to the younger sample. (**D-G**) Regenerating imaginal discs stained with an antibody that recognizes the Z1 isoform of the Broad transcription factor for wild-type (**D, E**) and *Ets21C* mutant (**F, G**) discs. Note that *Ets21C* expression was found to negatively correlate with *broad (br*) expression (**supplementary fig. 6**). Discs were dissected at the start of the regeneration period (0R) (**D, F**) and after 24h of regeneration (24R) (**E, G**). Note that BrZ1 levels increase from 0R to 24R and are at higher levels in the non-regenerating regions of the tissue and lower in the regenerating pouch region. Also note that *Ets21C* mutant discs have higher BrZ1 levels at 24R, especially in the region of the regenerating pouch. (**H-J**) To investigate the developmental progression of other tissues with the animal undergoing regeneration, we have counted the number of ommatidial rows within the eye-imaginal disc. Previous work has suggested that the growth of other imaginal discs may pause during the process of wing-imaginal disc regeneration, based on the overall size of the imaginal discs (Boulan *et al*., 2019). (**H**) Diagram of how the “eye-clock” can be used to compare the organismal-wide developmental progression. (**I**) The number of ommatidial rows for undamaged controls as compared to regenerating larvae. Note that the rate of addition of new rows of ommatidia added has slowed in the regenerating sample. (**J**) Comparison of the number of ommatidial rows for wild-type and *Ets21C* mutants during regeneration. Note that the *Ets21C* mutant animals show an increased number of ommatidial rows by 24h of regeneration, indicating that there is a reduction in the organismwide developmental delay that is observed in wild-type regenerating larvae.

**Supplementary figure 12.**
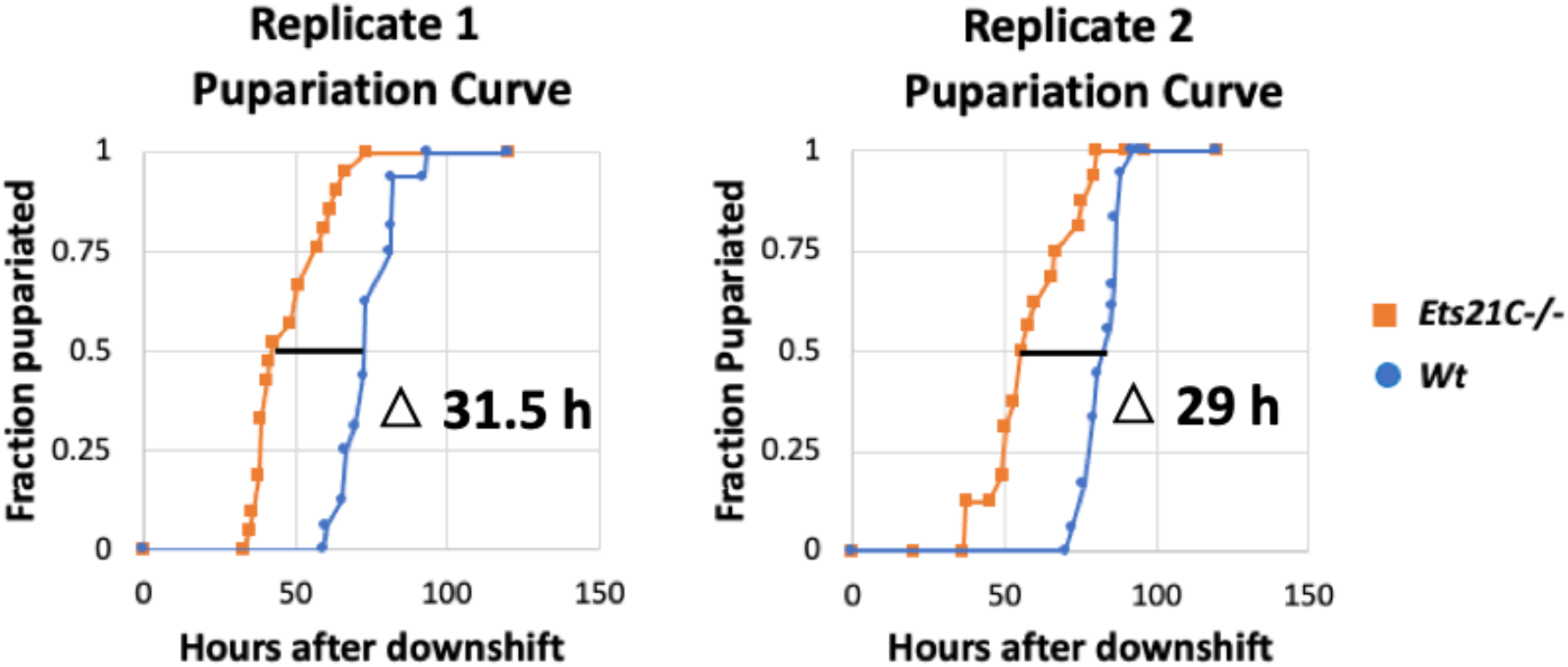
Relative pupariation timing for wild type (Wt) and *Ets21C* mutant animals following damage and regeneration. Images of vials were taken at 10-minute intervals and then scored based on coloration changes during pupariation. Replicates were biological replicates conducted on separate days. The relative difference in pupariation timing was calculated based on the point when one-half of the animals scored had pupariated. Following ablation and regeneration, *Ets21C* mutant larvae formed pupa 31.5 hours and 29 hours earlier than the wildtype controls.

**Supplementary figure 13.**
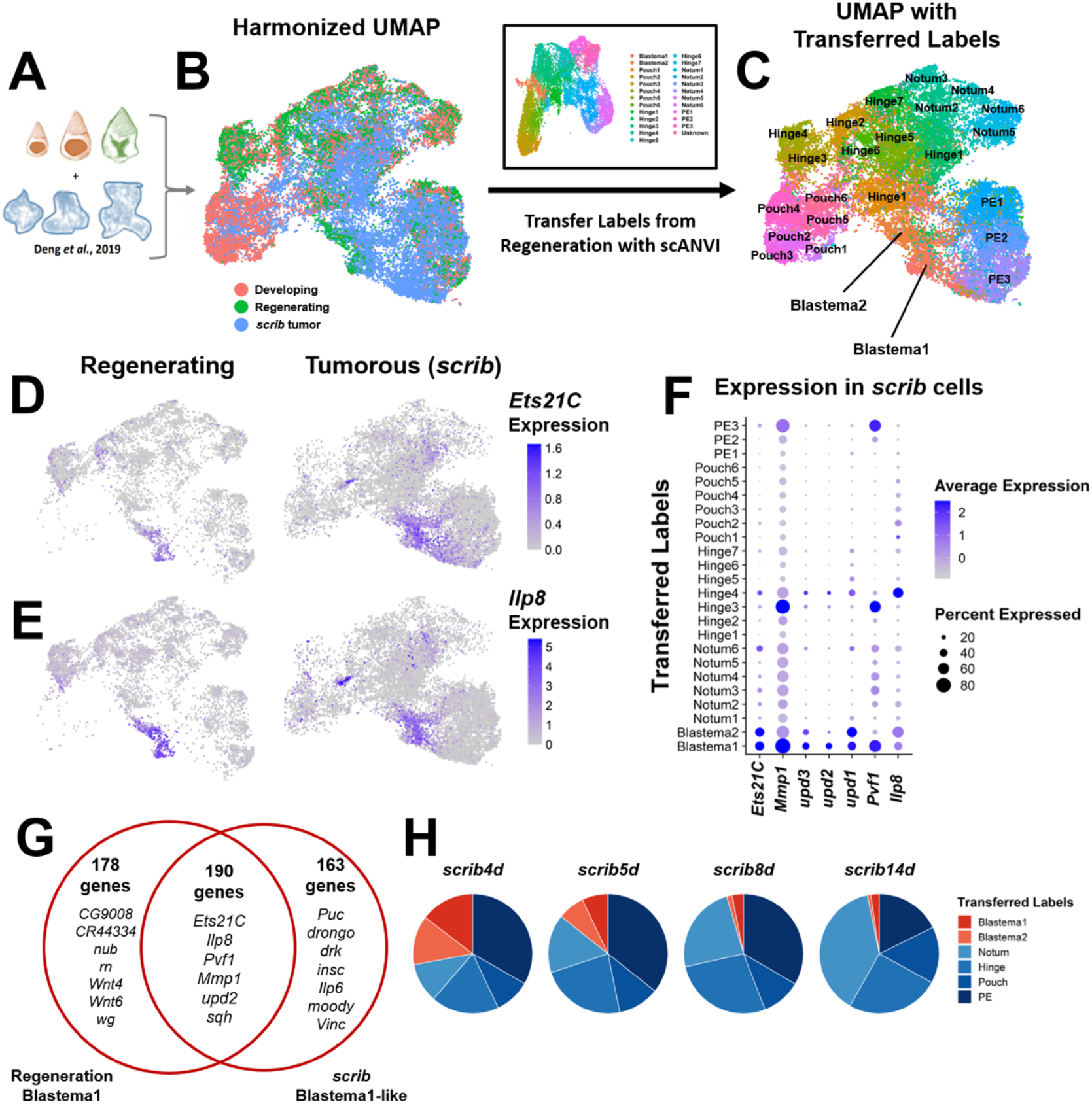
Single-cell comparison of regenerating and *scrib* tumorigenic discs. (**A**) Diagram of combined single-cell data. (**B**) Harmonized UMAP with cells colored by sample of origin (as also shown in **Figure 3Q**). (**C**) Harmonized UMAP colored by cell identity as generated by transferring labels from the regeneration cell atlas onto the *scrib* dataset using scANVI (see Materials and Methods). (**D, E**) Expression UMAPs of *Ets21C* (**D**) and *Ilp8* (**E**) in regenerating and *scrib* tumor datasets (note that expression for *scrib* tumors alone was also shown in **Figure 3R**). (**F**) Dot plot summarizing the expression of blastema markers within the *scrib* data. Note the co-expression of these genes in *scrib* clusters with Blastema1 and Blastema2 transferred labels. (**G**) Venn diagram comparing the overlap of markers for the Blastema1 cells within the regenerating data and the Blastema1-like cells within the *scrib* data. (**H**) Quantification of transferred labels within the *scrib* datasets, over the course of tumor development as collected by *Deng et al*. 2019 *(8)* (4d, 5d, 8d, and 14d = 4, 5, 8, and 14 days of tumor development, respectively). Note that there are more blastema-like cells on day 4 and 5 than on day 8 and 14 of within the *scrib* tumorous discs. This data is also summarized as “*scrib* early” and “*scrib* late” in **Figure 3S**.

## Notes

### Competing Interest Statement

The authors have declared no competing interest.

